# Proposed core role for cytosolic and transmembrane calpain cysteine proteases in mitotic cell divisions

**DOI:** 10.64898/2026.05.11.724225

**Authors:** Jennifer C. Fletcher, Mary A. Biggs, Hilde-Gunn Opsahl Sorteberg

## Abstract

Calpains constitute an ancient, extensive family of calcium-dependent cysteine proteases found in some bacteria and most eukaryotes. They are involved in a wide variety of developmental and cellular processes and are implicated in major human diseases, yet it remains to be seen if they have a common core function explaining their widespread and varied presence across taxa. Beyond their core CysPc catalytic domain, calpains contain diverse domain combinations and can be either cytosolic or membrane bound. Here we hypothesize a general role for both cytosolic and transmembrane calpains in cellular cytokinesis through positional anchoring and organization of microtubules (MTs). We propose that during plant cell division, the singular transmembrane calpain DEK1 localizes and organizes the array of cortical MTs from the microtubule organizing center (MTOC) to establish the location of the preprophase band and/or the site of cell plate formation according to the positional activation of DEK1 proteins in the nuclear membrane. Similarly, during cell division in animals, their calpains may be involved in setting the point of membrane invagination via their association with membrane-bound proteins. This proposition adds to the current picture of animal MTOC/centrosome function and suggests how a calcium peak during the initial cytokinetic furrowing might be transmitted. We discuss this novel mechanistic model for calpain activity in the context of data from the animal and plant literature, as well as of our novel discovery here of calpain sequences in both brown and red algal genomes. Finally, we speculate that the ancestral role of calpains in early eukaryotes, before the split into the major eukaryotic supergroups, may have been to facilitate the formation and function of MT arrays in flagella and cilia. From this origin, calpains may have developed new functions in eukaryote cell division processes by anchoring centrosomes/MTOC to set the cell division orientations that are especially important for complex multicellularity.

## Introduction

The growth and development of all multicellular organisms depend on cell divisions that are controlled by the cell cycle. Progress through the cell cycle is directed by cyclin-dependent-kinases (CDKs) that initiate DNA synthesis and drive the cell through the cycle. The mitotic cell cycle involves chromosome condensation and nuclear envelope breakdown, mitogen-activated protein kinase (MAPK)-mediated attachment of the chromosomes to mitotic spindle fibers, followed by sister chromatid separation and physical division of the cytoplasm and organelles by cytokinesis. A key facet of mitosis is the organization of microtubules (MT) into the mitotic spindle by the animal centrosome or plant microtubule organizing center (MTOC), cytoskeletal components that orient the chromosomes along a set division plane establishing the subsequent division and new cell wall position.

Several cell division mechanisms occur in eukaryotes. In animals, the cell division plane is determined by the position of the mitotic spindle anchored at the opposite ends of the cell by the centrosomes, consisting of two centrioles that organize the MT as the main MTOC. Cytokinesis then occurs through the formation of an actomyosin contractile ring that constricts the cell membrane to form a cleavage furrow that divides the cell in two across the short axis (Figure 1A). This ancient evolutionary process developed before the split of the eukaryotic supergroups (Pollard and O’Shaughnessy 2020; Yubuki and Leander 2013).

**Figure 1.**
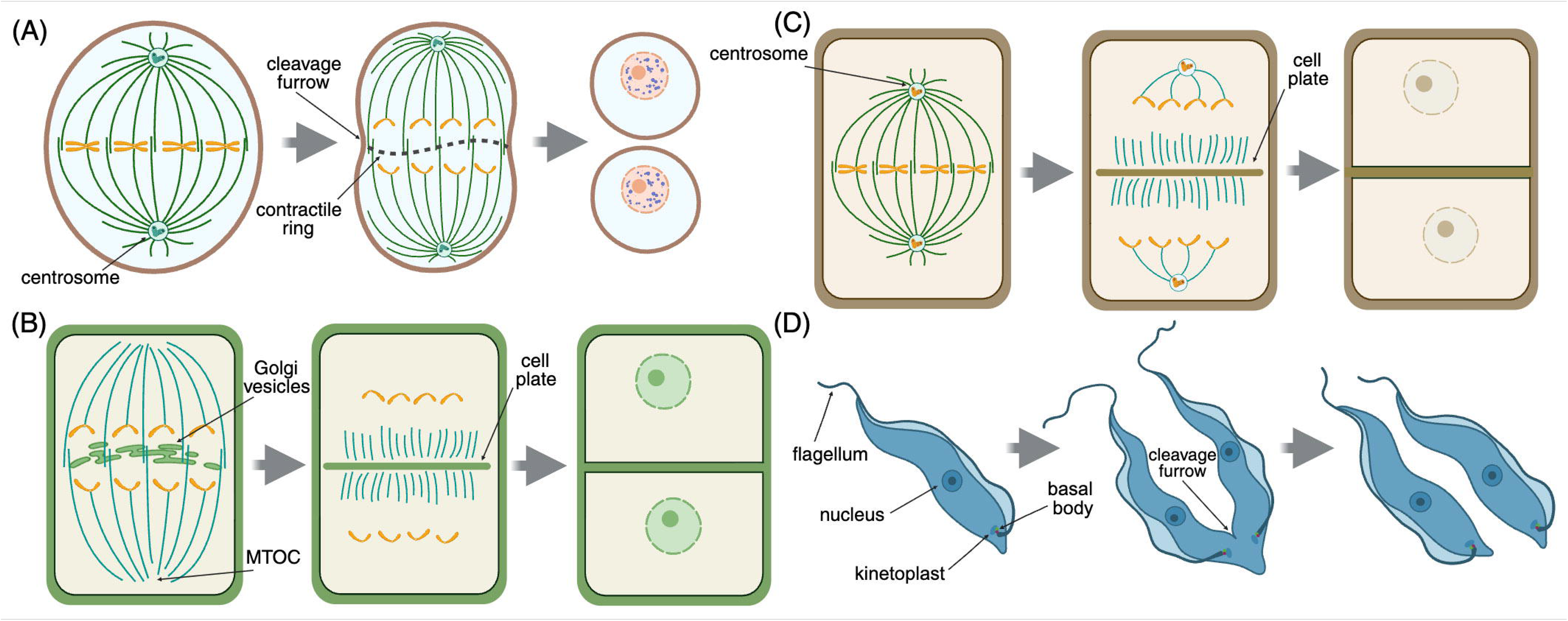
Schematics illustrating the process of cytokinesis in the somatic cells of various organisms. **(A)** Cytokinesis in animal cells. **(B)** Cytokinesis in land plant cells. **(C)** Cytokinesis in brown algal cells. **(D**) Cytokinesis in trypanosome cells. Figure generated using BioRender.

In most land plant cells, at the onset of mitosis the location of the cell division plane is set by the preprophase band (PPB). The PPB is a transient cortical ring of MT, actin filaments, endoplasmic reticulum, and associated proteins that is assembled and orientated perpendicular to acentrosomal MTOC activity and encircles the nucleus (Schaefer et al., 2017; Domozych and Bagdan 2022). The PPB leaves a positional cue at the future cell division site that guides the activity of phragmoplasts, assemblies of MT and Golgi vesicles that are organized by MTOC activity and fuse to form a cell plate. The cell plate then expands outward into a new cell wall that separates the daughter cells (Figure 1B). This group of organisms lack centrosomes and cell division depends on guidance by distributed MTOCs. Interestingly, the somatic cells of the basal liverwort plant *Marchantia polymorpha* initially form centrosome-like structures called polar organizers that appear before the PPB and serve as the MTOC (Buschmann et al., 2016). The cells subsequently divide like those of higher plants under direct guidance of the PPB by forming a cell plate (Brown and Lemon 2011). This dual mode of cytokinesis utilizing both centrosome-like structures and PPB is suggested to exemplify a stepwise evolutionary transition from green algae to land plants (Buschmann and Zachgo 2016) and is important to enable the asymmetric cell divisions that are especially important to meristem function, branching and setting 3D tissue orientation (Graham et al., 2000). Like land plants, most fungi also lack centrosomes, and their cell divisions are controlled by a MTOC named the spindle pole body (SPB) that plays a role in MT nucleation and additionally contributes to nuclear envelope breakdown (Jaspersen 2021).

Brown macroalgae, which are part of the SAR supergroup consisting of Stramenopiles, Alveolates and Rhizarians, have a combined set of structures and their cell division occurs via a combination of the two processes. That is, like animals, brown algae contain centrosomes that act as the MTOC (Pickett-Heaps,1974). Yet rather than forming a contractile ring and furrow they divide by centrifugal growth and Golgi vesicle-mediated fusion of wall material from inside to outside to form a cell plate (Nagasato and Motomura, 2002; Motomura et al., 2010), similar to land plants (Aoki et al., 2023) (Figure 1C). The latter behavior is likely explained by the fact that, like plants, brown algae harbor physiologically important cell walls allowing giant kelp species to grow impressive forest canopies, which depends on new cell wall deposition to complete cell division and grow large structures (Choi et al., 2024). Yet brown algae lack the cortical MTs, PPB and phragmoplasts present in land plants (Katsaros et al., 2006; Bellinger et al., 2023).

A fourth eukaryotic supergroup is the Excavata group of unicellular, mostly flagellated protists. Although it is debated whether this group is truly monophyletic (Burki et al., 2020), Excavates are generally characterized by their asymmetrical, “excavated” feeding grove and many undergo a specialized cell division by longitudinal fission where the parental cell splits lengthwise in two. This process is well described in the human parasite *Trypanosoma brucei*, in which basal bodies that anchor the flagella of the cell serve as centriole-like structures (Vaughan and Gull 2016). The basal body is a MT-composed organelle that like a centriole has a core ninefold-symmetrical MT and cartwheel structure and is physically connected to a kinetoplast consisting of mitochondrial DNA (Figure 1D). At the onset of mitosis, the basal body replicates and the new basal body assembles a new flagellum, the tip of which defines the position where cytokinesis initiates (Kohl et al., 2003). Cytokinesis then proceeds via ingression of a cleavage furrow along the longitudinal axis of the cell from anterior to posterior (Vaughan and Gull 2008; Wheeler et al., 2013; Zhou et al., 2016), without the formation of a contractile ring at the cleavage furrow site (Garcia-Salcedo et al., 2004). In all these cases, the positioning of the MTOC is critical for cytokinesis to proceed in the proper plane and for the correct partitioning of nuclei and organelles into the two daughter cells.

Calcium allowed life to emerge, especially multicellularity, and affects nearly every aspect of life by linking external stimuli to intracellular events via movement through ion channels or by release from cellular stores (Clapham 2007; Carafoli & Krebs 2016; Bootman & Bultynck 2020). Changes in calcium concentrations are universal regulators from bacteria to animals, evolutionary identified across early eukaryote linages before the split leading to the different phylogenetic groups of plants, brown algae, fungi and animals (Cai et al. 2014). Calcium plays a crucial role throughout the cell cycle in nuclear envelope breakdown, chromosome segregation, and cytokinesis (Nugues et al 2022). In fission yeast and animals, calcium spikes at the onset of cytokinesis initiate cleavage furrow ingression and subsequent plate formation (Atilla-Gokcumen et al 2010; Hepler 2015; Poddar et al 2021). At the molecular level, calcium is linked to the cell cycle by motility and motor proteins such as dynein and kinesin, RhoGTPases, initiation of the MAPK cascade and the cytoskeleton (Tan et al. 2006; Cai, 2014; Saternos et al 2020). Despite its pivotal role, a possible universal link remains to be detected between calcium and the many connected biological functions in most living organisms.

The transport of calcium into cells occurs via mechanosensitive channels, force-sensing integral membrane proteins found in all kingdoms of life that are activated by mechanical stimuli exerted on cell membranes (Ridone et al., 2019). Among these, Piezo plays an important role in regulating the cell cycle from centrosome integrity through divisions until cell death (David et al. 2022). In mammals, Piezo1 has been shown to anchor cytoskeletal components and functions upstream of calpains (Li et al., 2014; Friedrich et al., 2019; Zhang et al., 2021, Chuntharpursat-Bon et al 2023), Ca^2+^-activated cysteine proteases that are found in some bacteria and most all eukaryotes (Sorimachi et al. 2011). Mammalian calpains are grouped into classical and non- classical calpains based on their domain organization (Croall and Ersfeld 2007). The classical calpains contain a core CysPC (PC1 and PC2) domain as well as C2, calpain-type ß-sandwich (CBSW), and penta-EF hand (PEF) domains (Veselenviova et al 2022), whereas the non-classical calpains contain additional zinc finger (Zn), microtubule interacting and transport (MIT), and WW domain combinations (Ono and Sorimachi 2012; Safranek et al., 2023). All vertebrate and insect calpains are cytosolic proteins that function in the cytoplasm, not anchored to cell membranes, although they are associated with membranes and in some cases this association is linked to their activation (Araujo et al., 2018; Safranek et al., 2023). Calpains are central to many fundamental cellular processes, including cytoskeletal remodeling, cell signaling and apoptosis, centrosome functions, centrosome duplication and positioning, MT polarization, connecting chromosomes to spindles, chromosome positioning, and cell cycle progression (Honda et al. 2004; Storr et al. 2011; Valls et al. 2025). In mammals 14-15 genes encode calpain proteins, some of which are specific to certain tissues, whereas others are ubiquitous (Opsahl-Sorteberg et al., 2024).

In contrast, plants including primitive mosses and liverworts contain a single unique calpain called DEFECTIVE KERNEL1 (DEK1). Unlike animal calpains, at its amino terminus the DEK1 protein contains an extensive membrane anchor consisting of 21-24 predicted transmembrane domains interrupted by an unstructured loop or channel domain (Lid et al 2002; Kumar et al 2010; Perroud et al 2020). This is followed by an intracellular linker containing a LamininG-like domain (LGL), and the calpain protease comprised of a CysPC and a CBSW domain (Lid et al 2002; Johansen et al 2016). The DEK1 gene is highly expressed in actively dividing cells, including stem cells (Lid et al., 2005; Johnson et al., 2005; Liang et al. 2015), and the maize and Arabidopsis proteins are localized at the plasma membrane, ER and in endosome-like compartments (Tian et al. 2007; Johnson et al. 2008). During 3D growth in the moss *Physcomitrella patens*, PpDEK1 protein exhibits a more polarized localization restricted to the plasma membrane between recently divided cells (Perroud et al., 2020). Multiple studies have shown that DEK1 is essential for plant embryogenesis and post-embryonic development, epidermal identity, cell-to-cell adhesion, 3D cell orientation, stem cell maintenance and gene regulation, microtubule orientation, and cell wall positioning (Lid et al. 2002; Lid et al. 2005; Johnson et al. 2005; Tian et al. 2007; Johnson et al. 2008; Liang et al. 2015; Demko et al. 2024; Chen et al. 2024). These diverse biological roles have led previous investigators to conclude that DEK1 has a variety of activities within the cell (Demko et al., 2024). Here we hypothesize that *all* these activities can be explained by a single DEK1 function, and more generally, by shared calpain function in species that have multi-family calpains allowing for diversification into more specialized roles.

### Hypothesis for a primary role for calpains in cytokinesis

Here we present our hypothesis that calcium-regulated calpains play a fundamental role in cell mitosis, and likely meiosis, that has evolved with the diversification of cytokinesis mechanisms across eukaryotes (Table 1). In animals, cell division is mediated by the main centrosomal MTOC positioning that directs MT spindles, and then setting the position of the cytokinetic contractile ring forming the furrow that divides the cell. We propose that cytosolic calpains set the centrosome position, thereby directing and facilitating the progress of furrow ingression and enabling the completion of cell division at the proper position (Figure 2A). This activity may occur in response to calcium pulses that coincide with the initiation of furrow ingression (Atilla-Gokcumen et al 2010; Poddar et al. 2021) and may involve CysPC-mediated degradation of target proteins such as RhoA and fodrin. RhoA is a small GTPase that, when activated, stimulates actin nucleation and myosin activation to form the contractile ring and is sufficient for furrow initiation (Drechsel et al., 1997; Wagner and Glotzer, 2016). Fodrin is a non-erythroid form of spectrin that functions in metazoans to nucleate MT from centrosomes, elongate spindles, and position chromosomes at the metaphase plane, and is required for mitosis progression along with kinesins and dyneins (Shashikala et al 2013; Nellikka et al 2019; Sridhara and Shimamoto 2024). Additionally, cytosolic calpains may directly bind via their MIT and/or CBSW domain to the centrosomes in order to promote MT stability and position the plane of cytokinesis (Tonami et al., 2007).

**Figure 2.**
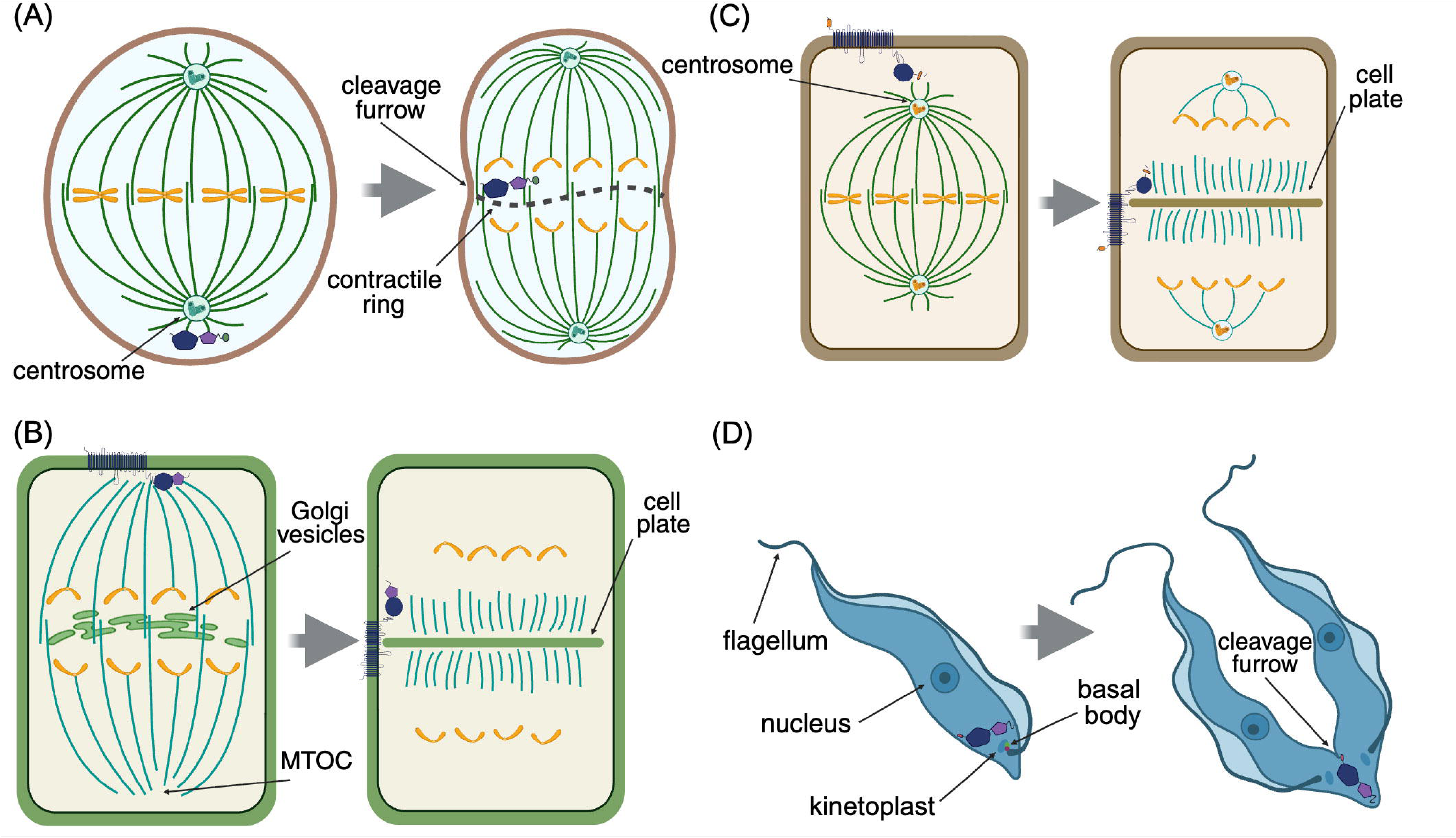
Schematics illustrating the proposed activity of calpains during mitosis in different types of organisms. **(A)** Cytosolic calpain function during cytokinesis in animal cells. **(B-C)** Transmembrane calpain function during cytokinesis in **(B)** land plant cells and **(C)** brown algal cells. **(D**) Cytosolic calpain function during cytokinesis in trypanosome cells. Figure generated using BioRender.

**Table 1.** Cytokinesis types, presence of centrosomes/microtubule organizing centers (MTOCs) and cilia/flagella linked to calpain domain combinations across the tree of life.

| Organism | Type of chromosome organizer | Cytokinesis type | Flagella/ Cilia movement | Additional calpain domains beyond CysPc<sup>1-2</sup> |
| --- | --- | --- | --- | --- |
| Cyanobacteria | NA | Fission | NA | ND |
| Dinoflagellate<br>*Symbiodinium minutum* | CL | Acrobase furrow | + | 44 calpain variants<br>ND |
| Trichomonas | CL attractophore | Ventral furrow | + | TML1 |
| Apusomad<br>Thecamonas | BB | Binary fission furrow | + | TML1&2, CBSW, MIT |
| Lower fungus<br>Yeast | SPB | Furrow | + | PalB |
| Fungus<br>*Aspergillus nidulans* | SPB | Septation cross walls | - | CBSW, MIT, PalB |
| Fungus<br>Magnaporthe<sup>5</sup> | SPB | Furrow | - | C2, EF |
| Parasitic Alveolata<br>SAR<br>*Plasmodium falciparum* | Centriolar plaque | Furrow | + | CBSW |
| Rhizaria SAR<br>Oomycete<br>*Phytophthora infestans* | + | Cleavage furrowing | + | TML3 & 4<sup>1</sup>, CBSW, MIT, C2, Zf-GRF |
| Ciliate Alveolata<br>SAR Tetrahymena<br>( ) | BB feeding + MTOC | Contractile ring | Cilia | TML1, TMS, EF |
| Stramenopila SAR<br>Brown algae (BA)<br>- | + | Centrifugal growth cell wall inside to out | + | TML, C2 |
| BA<br>*Desmarestia dudresnayi* | + | ND | + | TML, C2, EF |
| Red algae<br>*Rhodorus marinus* | Ring shaped MTOC | Furrow | - | Zf-GRF |
| Green algae (GA)<br>*Ostreococcus* | SPB | Phycoplast furrowing | - | Zf-GRF |
| GA<br>*Chlamydomonas* | BB/ MTOC | Phycoplast furrowing | + | CBSW |
| Unicellular biflagellate<br>charophyte GA<br>*Mesostigma viride* | BB | Centripetal cleavage | + | TML, CBSW |
| Basal land plants<br>Liverworts,<br>*Marchantia polymorpha* | PO | PPB | + | 2 TML variants,<br>CBSW |
| Higher land plants | MTOC | PPB+ | NP | TML2, CBSW |
| Animals<br>Opisthokonta | + | Furrow | Flagellum | C2, CBSW, MIT, PEF |
| CAPN7 most ancestral calpain | NA | NA | NA | MIT, PalBH |
| *Drosophila melanogaster* | + | Furrow | Flagellum | PEF, SOH |
| *Caenorhabditis elegans* | + | Furrow | Cilia | C2, MIT, PalBH, SOH |
| *Homo sapiens* | + | Furrow | Flagellum | MIT, CBSW, C2, PEF, PalBH |

In land plants, cytokinesis is initiated by two MTOC anchored at the opposite ends of the cell and their interconnected MT, which set the perpendicularly oriented cortical MTs into the PPB. The PPB positions the final cytokinetic division by the phragmoplast that directs cell plate formation and deposits cell wall material between the two newly forming daughter cells. The membrane-anchored calpain DEK1 is not required for PPB formation but is necessary for the arrangement of MTs into organized patterns and to correctly position the PPB (Liang et al., 2015). Therefore, we hypothesize that DEK1 may act to anchor the MTOCs and organize the subsequent positioning of the CMT and PPB, setting the new cell wall position and also facilitating the deposition of new cell wall material at the correct position by the phragmoplasts (Yan et al 2023).

Brown algal somatic cell mitosis depends on centrosomes that act as MTOCs, like animal cells, followed by plate formation to set the new cell wall, like land plant cells. Until recently it was not feasible to probe the calpain diaspora among the diverse families of brown algae, but the release of several dozen high-quality brown algal genomes (Diesel et al., 2023; Denoeud et al., 2024; De Weese et al., 2025) has enabled this type of investigation. We used the conserved CysPC domain amino acid sequence from *Arabidopsis thaliana* to identify calpain sequences across brown algal genomes. We identified a single amino acid sequence in each brown algal genome that aligns well with conserved CysPC domain amino acid sequences from oomycetes, land plants, moss and fungi (Figure 3). The CysPC domain consists of the PC1 (Figure 3A) and PC2 domains (Figure 3B) and contains three amino acids - cysteine, histidine and asparagine - that make up the catalytic triad in animal and land plant calpains (Berti and Storer, 1995; Hosfield et al., 1999). Among the brown algal CysPC sequences the asparagine residue in the PC2 domain is conserved among the majority of the brown algae species examined but has been substituted with a tyrosine residue in *F. serratus, P. canaliculata*, and *A. nodosum*, three members of the Fucaceae family (Figure 3B). The histidine residue at position 1848 in the PC2 domain is not conserved in any brown algal or oomycete species in our dataset, having been replaced with an alanine, serine or threonine residue (Figure 3B). Each brown algal and oomycete sequence instead features one or two adjacent histidine residues at positions 1781-82 in the PC2 domain (Figure 3B). The cysteine residue at position 1670 in the CysPC domain also is not present in any brown algae calpain sequence identified, having been substituted with a serine residue in many of them and by a threonine, alanine or glycine reside in the others (Figure 3A). This cysteine residue is likewise replaced by a serine, alanine or glycine residue in the PC1 domain of oomycete calpain sequences (Figure 3A). However, nearly all of the brown algal calpain sequences contain a cysteine residue at position 1632, and the oomycete calpain sequences contain a cysteine residue at position 1630 (Figure 3A). Taken together, these data indicate the presence of a well-conserved calpain CysPC domain in brown algal species that comprises both the PC1 and PC2 regions. However, the cysteine and histidine amino acids of the presumptive catalytic triad are found in different positions in the brown algae CysPC domains than they are in plants and animals; rather, their positioning closely resembles that found in oomycetes.

**Figure 3.**
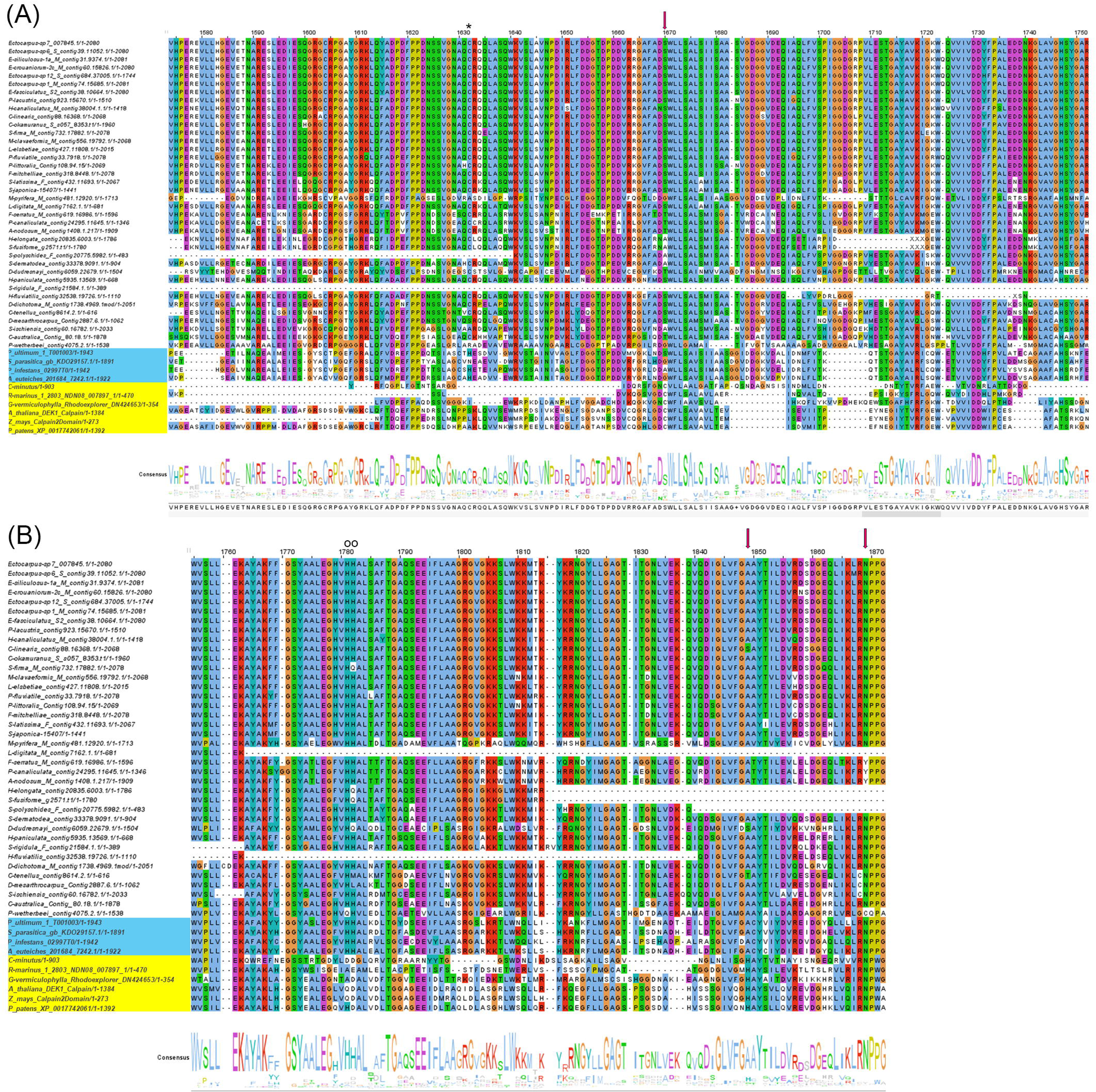
Multiple sequence alignment of the core CysPC domain of calpain sequences from brown and red algae as well as selected land plants and mosses. Alignment of **(A)** The PC1 subdomain and **(B)** the PC2 subdomain. Conserved amino acids are highlighted in colored boxes. Red arrows indicate the position of the amino acids in the catalytic triad (Cys-His-Asn). The red arrows indicate the positions of the three amino acids (C, H and N) that comprise the catalytic triad. The asterisk (*) indicates the position of the highly conserved cysteine residue in the brown algal sequences. Open circles indicate the positions of the highly conserved histidine residues in the brown algal and oomycete sequences. Oomycete species are highlighted in blue, and red algal, land plant and moss species are highlighted in yellow. The consensus amino acid sequence is shown along the bottom.

Most of the identified brown algal calpain sequences contain at least one other domain in addition to the protease core CysPC domain. However, in five brown algae genomes only a CysPC domain is predicted (Figure 4A), likely due to incomplete sequence information or annotation. The calpain sequences of two additional species, *D. mesarthrocarpus* and *H. paniculata*, are annotated as having a CysPC domain and a short C2 domain near the carboxyl terminus and may also be incomplete. We predicted the calpain sequences of *Saccharina latissima*, *S. japonica, S. promiscuus* and two other species based on several adjacent contigs (Figure 4A). The calpain sequences in these five and the majority of the other brown algal species are predicted to contain a series of TM domains animo terminal to the CysPC domain, and 18 species across several families are predicted to contain an additional C2 domain close to the amino terminus of the predicted protein (Figure 4A). Unexpectedly, the calpain sequence from *Desmarestia dudesnayi* uniquely harbors nested among the TM motifs two EF hand domains, which are present in some metazoan calpains but are absent from land plants (Opsahl-Sorteberg et al., 2024). Phylogenetic analysis of the brown algal calpain full-length amino acid sequences confirms the presence of a single calpain sequence in each brown algae genome (Figure 4B), as well as strong sequence conservation within the various brown algal families represented. We hypothesize that the single TM calpain is utilized during cell division in brown algal cells to set the position of the centrosome, similar to the situation in animal cells (Figure 2C).

**Figure 4.**
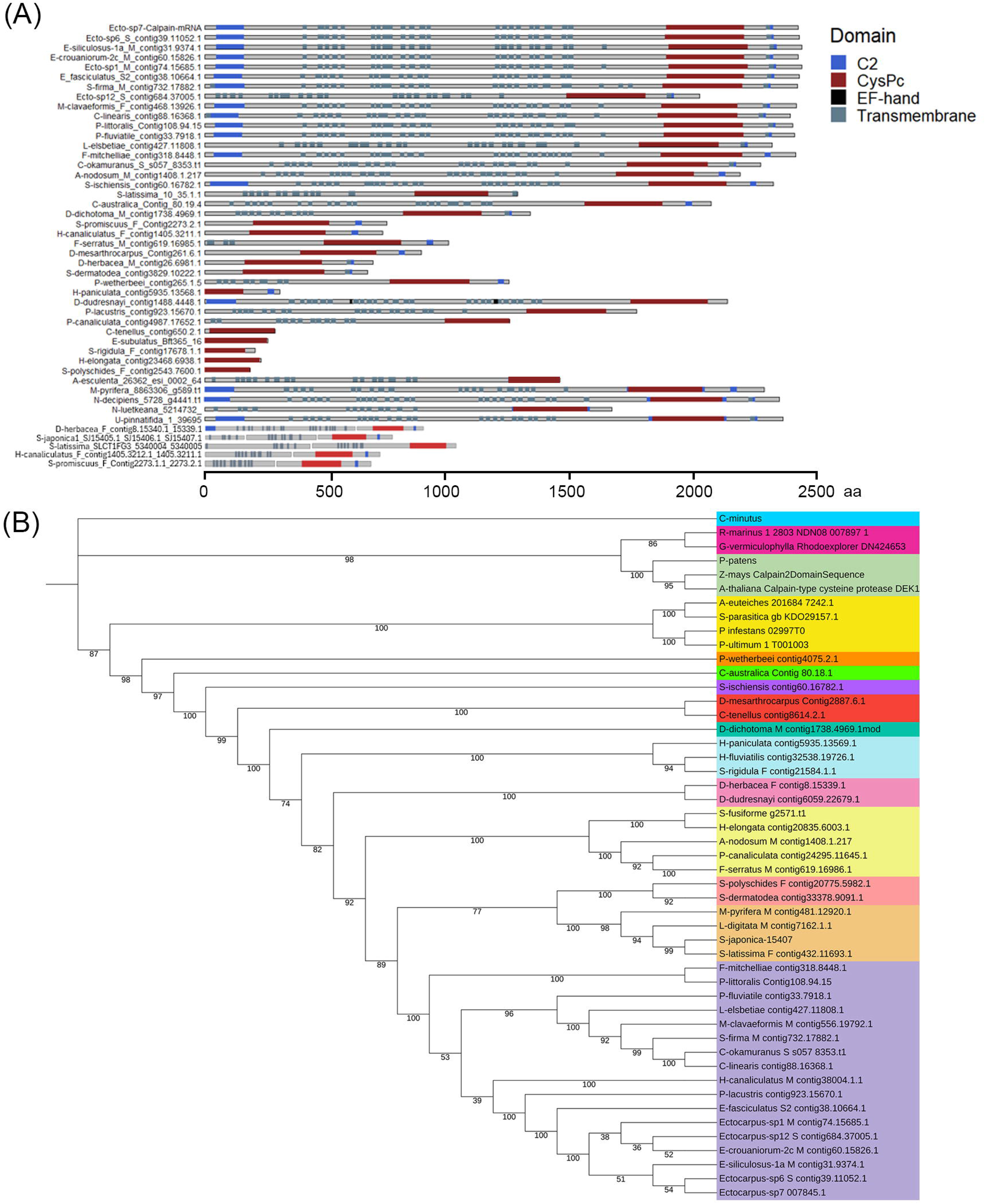
Calpain sequences in brown algae. **(A)** Domain composition of brown algae calpain amino acid sequences. The calpain sequences in five of the brown algae are discontinuous and are shown with the corresponding sequence gaps. **(B)** Phylogenetic tree of the brown algae calpain sequences. The tree was constructed using the IQtree program embedded in PhyloSuite. Bootstrap values on each node indicate the proportion of recovered nodes out of 1,000 bootstrap replicates. Bright blue indicates *C. minutus*, purple indicates red algal species, pale green indicates land plant and moss species, yellow indicates oomycete sequences, and dark blue indicates *S. ischiensis*. The other colors indicate species within the same brown algal family.

Beyond the calpain sequences we characterized in the brown algal genomes, our query of red algal genomes and transcriptomes using the conserved CysPC domain amino acid sequence from *A. thaliana* returned a hit from *Rhodosorus marinus* and one from *Gracilaria vermiculophylla*. The CysPC domain of both red algal calpain sequences contains the Cys-His-Asn catalytic triad in the same positions as in the green algal and land plant CysPC domains (Figure 3). Overall, the predicted *Rhodosorus marinus* calpain protein features a GRF zinc finger Zf_GRF-CycPC domain configuration also present in green algae, phytoplankton, and all *Plasmodium* species within the Aveolates (Russo et al., 2009; Zhao et al., 2012). The predicted *Gracilaria vermiculophylla* calpain protein features a CysPC-CBSW domain architecture present in nearly all eukaryotic supergroups and considered to be one of the ancient calpain types (Zhao et al., 2012). These findings indicate that calpains are present across the red and green algal and land plant lineages of the Archaeplastida supergroup.

Our hypothesis that calpains play a primary role in cell mitosis by positioning the MTOC and the cleavage furrow or PPB and thereby orienting the cell division orientation is consistent with known functions for these proteins in the animal cell division process. Cytosolic calpains already have been shown to function in MT- associated processes during mammalian mitosis. In human HeLa cells, CAPN2 protein levels are elevated during mitosis, and this activation is necessary for proper MT attachment to kinetochores and correct alignment of the chromosomes at the spindle pole prior to their segregation into the two daughter cells (Honda et al., 2004). In human breast cancer cell lines, CAPN2 plays a key role during the progression of mitosis and cytokinesis by reducing the levels of the LIM Kinase-1 (LIMK1) protein that phosphorylates the actin-severing protein cofilin-1 (CFL1) at critical steps of mitosis (Rodriguez-Fernandez et al., 2021). A similar mechanistic role for calpains in indirectly modulating the activity of MT binding proteins during mitosis can likewise be envisioned. CAPN2 is also known to degrade fodrin (Siman et al., 1984; Sato et al., 2004), which in brain cells co-localizes with MTs and plays a role in MT spindle organization and mitotic progression by associating with γ-tubulin to promote the transport of the tubulin to centrosomes for MT nucleation (Shashikala et al., 2013; Nellikka et al 2019). Fodrin concentrations peak prior to the initiation of mitosis, and then the proteins dissociate from the centrosomes after prophase (Shashikala et al., 2013). It has been hypothesized that calpains mediate the cleavage of fodrin from γ-tubulin to ensure the proper progression of mitosis (Shashikala et al., 2013). Thus, fodrin may be a key target of cytosolic calpain activity during mitotic microtubule nucleation in metazoans, where it is predominantly found (Sreeja et al 2022).

In addition, some non-canonical mammalian calpains contain a microtubule interacting and trafficking (MIT) domain that is absent from calpains found thus far in bacteria and tetrahymena (Zhao et al., 2012; Opsahl-Sorteberg et al. 2024). The MIT domain forms an asymmetric three-helix bundle structure that resembles the first three helices in a tetratricopeptide repeat (TPR) motif (Scott et al., 2005). The MIT domain can bind to ESCRT-III proteins that constrict membranes and mediate fission, including during cytokinesis where they mediate abscission of the dense central array of MTs called the midbody to create the two daughter cells (Wenzel et al. 2022). During cytokinesis in humans, the ESCRT-III subunit IST1 recruits CAPN7 proteins to midbodies, where its proteolytic activity is required for abscission checkpoint maintenance and at the final stage of cell division to complete the abscission process (Wenzel et al. 2022; Paine et al. 2023).

CAPN6 is another non-canonical mammalian calpain, one that lacks an MIT domain as well as the active-site cysteine residues necessary for protease activity (Dear et al., 1997). It functions as a microtubule-stabilizing protein that associates with microtubules via its CBSW domain and colocalizes to microtubule structures, including the central spindle and midbody, during cytokinesis (Tonami et al., 2007). During mitosis, CAPN6 protein is associated with the mitotic spindle and appears to be required for the progression and completion of cytokinesis, similar to fodrin, as its over-expression retards the ingression of the cleavage furrow in human cells (Tonami et al., 2007; Nellikka et al 2019).

In contrast to the body of understanding of cytosolic calpain activity during the animal cell cycle, nothing is known about the cellular mechanism of DEK1 calpain function. DEK1 is localized to the plasma membrane as well as internal membranes (Tian et al., 2007; Johnson et al., 2008), and regulates cell wall composition and structure (Demko et al., 2014; Galletti et al., 2015; Amanda et al., 2016; Amanda et al., 2017), including the synthesis of cellulose and pectin and the mechanical properties of the primary cell walls (Novakovic et al, 2023). Further, DEK1 is suggested to facilitates mechanosensitive calcium transport through the association of its TM domains with a yet unidentified rapid mechanically activated (RMA) channel in the plasma membrane (Tran et al., 2017). This RMA channel displays electrophysiological properties similar to those of mouse Piezo channels (Coste et al., 2010) and may be encoded by a possible unidentified plant PIEZO gene or a different protein with a similar function. Piezo has 24 TML domains, comparable to DEK1’s 23, and the two proteins have seemingly shared functions (Guerringue et al., 2018). Piezo1 and 2 localize to the centrosome and play roles in cell cycle progression (David et al., 2022) and additionally are localized to the cleavage furrow controlling final cell division (Carrillo-Garcia et al., 2021). We propose that perception in the plasma membrane of mechanostimuli from the internal tension of the dividing cell leads to transiently elevated calcium transport either through the DEK1 loop/channel structure, which itself shows properties of mechanosensory channels (Guerringue et al., 2018; Yan et al 2023), or via its association with a separate RMA channel such as Piezo1. This causes an increase in cytosolic calcium concentration that triggers DEK1 auto-activation (Johnson et al., 2008) and release of the activity to affect the cytoskeleton positioning the MTOC and orient the new cell wall. Considering that the protease-deficient human CAPN6 calpain co-localizes to microtubule bundles via its CBSW domain and promotes their formation and stabilization (Tonami et al, 2007), we propose DEK1 may act through its CBSW domain to directly bind and stabilize the MTOC to set the new cell division plane.

Our hypothesis is consistent with multiple lines of evidence regarding DEK1 activity within cells. First, although localized to the plasma membrane, ER and endosome-like structures, DEK1 signaling is cell autonomous (Becraft 2000). Second, DEK1 affects cell wall orientation, where the PPB and cell walls are incorrectly positioned in severely affected Arabidopsis *dek1* early embryos (Liang et al., 2015). Third, the MTs in *dek1* protoderm-like cells are orientated more randomly than in wild-type cells (Liang et al., 2015). An altered cortical MT arrangement was also documented in the epidermal cells of partial loss-of-function *dekl-4* plants (Galletti et al., 2015). These data indicate that DEK1 is important for the proper arrangement of the cortical microtubule systems, which in turn may affect the orientation of the phragmoplasts during mitosis and subsequent cell wall deposition. Several studies in Arabidopsis and moss consistently indicate a role for DEK1 in position-dependent cell wall orientation (Demko et al., 2014; Perroud et al., 2014; Liang et al., 2015). In addition, the expression of multiple genes related to the MTOC/centrosome and nuclear envelope degradation (NEBD) is altered in Arabidopsis *dek1* mutants or over-expression lines, such as Human CENP-E and animal KID, MAP65, mitotic kinesin, NEK5, CDKs, RanGAP, E2F and ROPs (Liang et al., 2015). Many of these genes play prominent roles in regulating phragmoplast MT dynamics during cytokinesis (Smertenko et al., 2018). Similarly, moss DEK1 target genes include those involved in reorientation of the phragmoplast and the cell division plane (Demko et al., 2024). PpDEK1-regulated transcription factors are significantly enriched for NERD-like calpain cleavage patterns, which may direct them to the N-end rule degradation (NERD) pathway for subsequent ubiquitin-mediated proteolytic degradation (Demko et al., 2024), analogously to the targets of mammalian calpains (Piatkov et al., 2014). We propose that DEK1 may directly sense and respond to mechanical stresses during the cell division process and transform these mechanical stimuli into calcium-based signals, which then trigger a downstream signal transduction pathway regulating the expression cytokinesis-related genes, similar to the mechanism proposed by Yan et al., 2023.

Although direct targets of DEKI protease activity have yet to be identified, potential substrates during mitosis include NAC WITH TRANSMEMBRANE MOTIF (NTM1), a membrane-bound NAC domain transcription factor that is activated by proteolytic cleavage to mediate signaling by the plant hormone cytokinin during cell division in Arabidopsis (Kim et al., 2006). DEK1 also has been proposed to affect cellulose synthase trafficking and mobility in the plasma membrane, either directly by regulating CELLULOSE SYNTHASE (CESA) complexes (CSCs) at the post-translational level or indirectly through interactions with CESA regulatory proteins or cytoskeletal components that guide CSCs (Safranec et al., 2023). Candidate factors include the endo-1,4-beta-glucanase KORRIGAN (KOR) that is involved in CESA regulation (Mansori et al., 2014; Vain et al., 2014), CSII/POM2 that facilitates binding between CSCs and cortical MTs (Gu et al., 2010; Bringmann et al., 2012), and PATROLI (PTL1) that interacts with CS1/POM2 and exocyst complex proteins to deliver CSCs to the plasma membrane (Zhu et al., 2018).

## Discussion

Calpains are calcium-dependent cysteine proteases crucial to eukaryotic cellular processes including various aspects of cell cycle activity. A vast array of calpain architectural combinations exist in eukaryotes, likely originating from the domain shuffling of four combined variants early during eukaryotic evolution (Zhao et al, 2012). These domains were present in eubacteria and archaea before being combined with the core CysPC domain that defines a calpain based on its catalytic activity. Extant unicellular protists and streptophytes contain long membrane-anchored calpains, and the ciliate *Tetrahymena* genome encodes up to an impressive 26 calpain genes (Croall and Ersfeld 2007), suggesting an ancient functional role important to these singled celled organisms. Here we hypothesize that the core role of calpains is to set the position of the MTOC during mitosis, and possibly meiosis as well. We propose that during cytokinesis in plants, the transmembrane TML calpain DEK1 anchors the MTs in the MTOC relative to the cell membrane to set the cell division plane and orient the subsequent phragmoplast-mediated formation of new cell walls. Similarly, during cytokinesis in animals the centrosome, a type of MTOC, initiates furrowing perpendicular to the centrosome axis of MT by combining a calcium pulse detected prior to furrowing with the activation of the cytosolic calpain CysPC domain that then cleaves multiple substrates and anchors and/or stabilizes the MT bundles, which allows open chromosome separation and the allocation of organelles into the daughter cells.

Mitosis in brown algae occurs through a combination of the MTOC-driven processes observed in plants and animals, suggesting that brown algal cytokinesis might involve calpain activity. Previous work revealed the presence of a single DEK1-like TM calpain sequence in the reference genomes of 17 brown algae species spanning multiple families (Denoeud et al., 2024). Our study expands on and provides a more detailed analysis of these putative membrane-bound calpain sequences. We demonstrate that the genomes of the more than three dozen brown algal species investigated each contain a highly conserved CysPC domain sequence consisting of both the PC1 and PC2 regions (Figure 3), although the domains in some species are incomplete likely due to lower genome sequence quality. Interestingly, the positioning of the Cys-His-Asn conserved active site residues comprising the catalytic triad is not conserved in the brown algal calpain sequences. Instead, all but three of the brown algal calpain sequences contain an asparagine residue at the conserved position at the end of PC2, but have a cysteine residue at a more amino-terminal position in PC1 (Figure 3A) and one or two histidine residues at a more amino-terminal position in PC2 (Figure 3B). A parsimonious interpretation of our data is that these calpains may lack protease activity, similarly to the mammalian CAPN6 protein (Matena et al 1998; Tonami et al., 2013). However, some animal calpains with substitutions in these residues do not display loss-of-function phenotypes (Spadoni et al., 2003), suggesting either that protease activity is not completely abrogated or that these calpains may have functions beyond proteolysis, such as in signal transduction and/or gene regulation. On the other hand, we note that the positions of the cysteine and histidine residues in the CysPC domains of brown algae calpains are nearly identical to those in oomycetes (Figure 3). The conservation of spacing of these key amino acids within these Stramenopile lineages that shared a last common ancestor over 400 million years ago may indicate that the brown algal and oomycete calpains contain a functional but non-canonical catalytic triad. Further biochemical experiments will be required to determine if this non-canonical triad retains protease activity *in vivo*.

Analysis of the domain architecture of brown algal calpain sequences revealed that each contains a CysPC domain consisting of PC1 and PC2 along with one or more C2 domains and several stretches of multiple TM domains, as many as 26 in total. The putative calpain sequences identified in a few brown algal species currently lack TM or C2 motifs. However, because only a single CysPC core calpain domain sequence is present in any brown algal genome (Figure 3 and 4), the most parsimonious interpretation of these data is that the sequences are incomplete and that the full domain configuration is likely to be C2-TML-CysPC-C2. Although C2 domains are present in classical animal calpains and TM domains are present in land plant calpains, the brown algal calpain C2-TML-CysPC-C2 domain configuration is so far unique among eukaryotes (Zhao et al., 2012; Safranek et al., 2023). This architecture represents a fifth type of eukaryotic membrane-anchored calpain, most similar to the Type 3 C2-TML-Linker-CysPc-WW version found in various oomycetes (Safranek et al., 2023), which like brown algae are members of the Stramenopiles subclade. The predicted *D. dudesnayi* C2-TML-EF-TML-EF-TML-CysPC domain structure is also unique, among not only the brown algae but eukaryotes in general, although an EF-CysPC configuration is present in *Tetrahymena thermophila* (Zhao et al., 2012), an Alveolate member of the SAR supergroup. Further investigation will be required to discern whether these *D. dudesnayi* calpain putative EF hand motifs are functional and what their evolutionary origin might be.

Our characterization of calpain sequences in brown algae emphasizes the modularity of these multifaceted cysteine proteases across eukaryotes, and their specific domain combinations hint at possible evolutionary shared cellular functions. Beyond the CysPC protease core domain, the C2 domain primarily acts as a calcium-binding motif that is involved in signal transduction and membrane association (Stahelin and Cho 2001; Bondada et al., 2021; Safranek et al., 2023). Through the combination of the C2 and TM domains, these brown algal calpains therefore potentially have the capacity to detect and transmit calcium signals across the plasma membrane. Further, based on their overall sequence similarity to *A. thaliana* DEK1, the brown algal calpains may bind and stabilize MTs in their centrosomes during cytokinesis, and this may be a primary function for calpain across the eukaryotic supergroups. It is also worth noting that our analysis likely underestimates the true number of brown algal calpains, because we manually removed sequences that lacked a full length CysPC domain even if they displayed significant amino acid alignment. More complete sequencing and annotation of brown algal genomes will provide a more comprehensive picture of the nature of the calpain family in this important group of marine Stramenopiles.

The centriole is an ancient MT-based cylindrical organelle that not only nucleates the formation of the centrosome, the main MT organizing center in animal cells, but also can be modified to form basal bodies that template the formation of cilia and flagella. The centrosome templates the formation of the primary cilium from one of its centrioles, which is then assembled at the plasma membrane (Joukov & De Nicolo 2019). There is a direct structural relationship between MT and membrane furrowing during ciliogenesis, where cilia initiate as a basal membrane invagination/ pocket, via interaction with actin-regulating proteins, Rho GTPases and formins (Birkenfeld et al., 2008; Paridaen et al., 2013; Onishi et al., 2020; Wilsch-Bräuninger & Huttner 2021). In fact, the presence of centrioles correlates across the tree of life with the presence of cilia but not of centrosomes, suggesting that the ancestral role of centrioles was to direct cilia formation and activity (Carvalho-Santos et al., 2011; Breslow and Holland, 2019). Thus, during evolution the centriole likely derived from a basal structure that nucleated MTs in flagella and cilia in the LECA and was subsequently lost in land plants.

Calcium signaling is involved in a variety of cilia-dependent biological processes, including mechanosensation, chemosensation, the cell cycle, cell polarity and cell migration (Saternos et al., 2020), and evidence exists for a role for calcium-dependent calpains in mammalian cilia and flagella. Calpains are colocalized with cilia components in multiple organisms including in the green algae *Chlamydomonas reinhardtii* flagellar proteome, in Trypanosomes associated with cilia functions as calpains are essential for flagellar attachment/structure, in sea urchins for mediating sperm activation, and in human sperm (Ortiz-Garcia et al., 2023). In mouse fibroblast cells, CAPN6 acts as an inducer primary of ciliogenesis that is proposed to increase the levels of alpha-tubulin as a post-translational modification to promote MT stability and function (Kim et al., 2019). In spermatozoa, which use flagella for motility, calpain activity is essential for physiological processes such as capacitation, acrosomal reaction, mobility, and fusion that are required for successful fertilization (Rojas and Moretti-Rojas, 2000; Aoyama et al., 2003; Ozaki et al., 2001; Ashizawa et al., 2006). Specifically, CAPN1 regulates the remodeling of the spectrin cytoskeleton (Bastian et al., 2010) as well as lipid raft rearrangement and activation of the Src kinase family (Maldonado-Garcia et al., 2017). Membrane contractile networks are especially fundamental to cytokinesis and cell motility, depending on the cell structures actin, tubulin and MT and their crosslinking to proteins and membranes. Common to these functions are MT organized in centrosomes and our hypothesis is that these functions may all be controlled by calpains. We speculate that early during eukaryotic evolution, calpains may have developed roles in cytoskeletal functions involving MT regulation during cilia/flagella formation and regulating MTOC and cell wall positioning activities during cytokinesis.

Our hypothesis both raises new questions and opens novel avenues for the investigation of calpain function at the cellular and biochemical levels. One key question is how the cytosolic calpains set the position of the cell division plane when they themselves are not membrane anchored. The answer may lie in their association with membrane-bound proteins that have a specific sub-cellular localization, but this remains to be determined. The sub-cellular localization of TML calpains during mitosis in different cell types also warrants additional careful investigation. In addition, the proteolytic substrates of transmembrane calpains such as DEK1 are as yet unknown, and whether such TML calpains can directly bind MTs needs to be tested. The relative contribution of calpain proteolytic activity and MT anchoring activity towards facilitating MTOC orientation during cell division also remains to be assessed. We hope this hypothesis and theory paper will provide a useful foundation for further studies of this diverse family of key regulatory proteins for the future benefit of agriculture, aquaculture, and the treatment of calpain-mediated human diseases including dementia and cancer.

## Methods

### Brown Algae Genome Mining for Calpain Sequences

To identify putative DEK1-like calpains in brown algae, the Phaeoexplorer brown algae protein database (Denoeud et al., 2024) was queried using BLASTp with a DEK1-like calpain sequence from *Ectocarpus* sp. 7. The domain compositions of the top 100 hits from this search were predicted individually using InterProScan (Jones et al., 2014), and the output filtered to exclude sequences lacking a predicted CysPC domain. To visualize and compare the domain compositions of the putative brown algal calpains, this filtered output file was uploaded to RStudio. Additionally, the 10 available brown algal proteomes and 26 red algal proteomes in the PhycoCosm database (Grigoriev et al., 2021) were individually queried using BLASTp with the same DEK1-like sequence from *Ectocarpus* sp. 7. The domain compositions of the top hits from these queries were predicted using InterProScan. The output was filtered as described, and the remaining sequences uploaded to RStudio.

To generate the brown algal calpain domain composition schematic in RStudio, InterProScan domain prediction outputs were first manually reformatted in Excel for compatibility with the drawProteins package (Brennan, 2018). Extraneous annotations and structural descriptions were filtered out, and entry names were optimized for clarity. The ggplot2 package was then used to construct the initial plot, followed by domain architecture visualization using drawProteins. The BiocStyle and knitr packages in RStudio were employed to visually optimize and format the figure.

### Multiple Sequence Alignment and Phylogenetic Analysis

To probe the composition of the brown algal CysPC domain, a sequence alignment was generated using canonical CysPC domains from plants and cyanobacteria alongside selected DEK1-like sequences from heterokonts, including brown algae, as identified in Denoeud et al., 2024. Cyanobacterial sequences were obtained from Veselenyiova et al., 2022, and the *Physcomitrella patens* sequence (Johansen et al., 2016) was downloaded from the reference genome in NCBI GenBank (Rensing et al., 2008). *Arabidopsis thaliana* and *Zea mays* sequences were downloaded from Uniprot (Theologis et al., 2000; Lid et al., 2002; Yi et al., 2011). Sequences were aligned by Multiple Alignment using Fast Fourier Transform (MAFFT), followed by automatic trimming using TrimAI (Capella-Gutiérrez et al., 2009; Katoh and Standley, 2013). Following the generation of a trimmed alignment file, the sequences were visualized using JalView (Waterhouse et al., 2009).

To generate the phylogenetic tree, the trimmed alignment file was filtered to include only the heterokont sequences from the Phaeoexplorer database. This alignment file was then loaded into PhyloSuite v2.0dev2 and a treefile generated using the IQ-TREE plugin with *Schizocladia ischiensis* included as the outgroup (Zhao et al., 2025; Zhang et al., 2019; Xiang et al., 2023). Bootstrap values were calculated with 1000 replicates. The resulting treefile was uploaded to the ITOL webserver (Letunic and Bork, 2024) where the final phylogenetic tree was visualized and formatted. Entry names were optimized for clarity.

## Acknowledgements

We are grateful to all the authors of published work that have led us to this synthesis and hypothesis. Special thanks to Huw Jones for many walks talking through results leading to the equatorial plane and cell wall positioning by DEK1. Thanks to Mark Cock, Sheila McCormick and Odd-Arne Olsen for bringing us together and leading the way with brown algae and calpains, Sebastian Svensson for helpful discussions on the manuscript, and Martin Paliocha for improving our understanding of parallel evolution.

## Author Contributions

H-G O-S and JCF conceived the article subject and wrote the manuscript. MAB generated the new data, generated figures and commented on the drafts. All authors accepted the final version of the manuscript.

## Funding

H.-G.O.-S. was funded by the Norwegian University of Life Sciences faculty of Biosciences and M.A.B. was funded by the University of California, Berkeley Department of Plant and Microbial Biology. J.C.F. was funded by the United States Department of Agriculture (CRIS 2030-21210-001-00D) and the United States National Science Foundation (NSF-OISE 2435380).

## Conflict of Interest

The authors declare that the research was conducted in the absence of any commercial or financial relationships that could be construed as a potential conflict of interest.

## Notes

### Competing Interest Statement

The authors have declared no competing interest.

